# The microbiome of captive hamadryas baboon

**DOI:** 10.1101/2020.01.10.901256

**Authors:** Xuanji Li, Urvish Trivedi, Asker Daniel Brejnrod, Gisle Vestergaard, Martin Steen Mortensen, Mads Frost Bertelsen, Søren Johannes Sørensen

## Abstract

Hamadryas baboon is a highly social primate that lives in complex multilevel societies exhibiting a wide range of group behaviors akin to humans. Here, we report the comprehensive 16s rRNA gene analyses of group-living baboon microbiota across different body sites. Additionally, we compared the baboon and human microbiome of the oral cavity, gut and vagina. Our analyses show that the baboon microbiome is distinct from the human and baboon cohabitants share similar microbial profiles in multiple body sites. The oral, gut and vagina shared more bacterial ASVs in group-living baboons than in humans. The shared ASVs in baboons had significantly positive correlations, suggesting a potential bacterial exchange throughout the body. No significant differences in baboon gut microbiome composition within the maternity line and between maternity lines were detected, suggesting that the offspring acquire their gut microbiota primarily through bacterial exchange among cohabitants. Besides, *Lactobacillus* was not so predominant in baboon vagina as in the human vagina but was the most abundant genus in baboon gut. These data and findings can form the basis of future microbiome studies in baboons and be used as a reference to research where the microbiome is expected to impact human modeling with baboons.

## Introduction

Humans and other primates are home to trillions of symbiotic microorganisms. Interactions between a host and its microbes affect host physiology, behavior, reproduction, immunity and evolution [1–3]. The Human Microbiome Project, through monitoring or manipulations of the human microbiome, helps us better understand the associations between microbes and human health [4]. In contrast to the widely studied human microbiome, there is a paucity of information on the host-associated microbiomes of nonhuman primates (NHPs). Information about NHP microbiota is essential for understanding the factors underlying microbial coevolution with their hosts [5, 6]. Broad primate microbiome surveys could also allow for the development of predictive biomarkers to improve nonhuman primate health and management.

Baboons are one of the most biologically relevant research animal models for humans due to their genetic and physiological similarities to humans [7]. Baboons (genus *Papio*) are large-bodied, omnivorous, highly social, terrestrial Old World African monkeys that occupy a wide array of habitats similar to those of early hominins [8, 9]. Of the six recognized species [10], the social system of hamadryas baboons in particular shares even more similarities with humans than that of other baboons [9]. Like modern humans, the hierarchical social networks of hamadryas baboons connect individuals at multiple levels [8]. Frequent social interactions (mostly grooming) are necessary for baboons to maintain affiliative bonds [11].

To our knowledge, only the rectal and vaginal microbiota of baboons have been examined, likely because baboons can be used as a model in female reproductive studies due to the similarities shared with human reproductive tracts [12–14]. Investigations into human lung microbiology are a relatively new field [15], largely because of technical hurdles, ethical considerations and small sample sizes [16], whereas the use of baboon models can provide novel information useful in investigations into the pulmonary microbiome. In this study, we investigated 184 samples of rectal, oral, oropharyngeal, cervical, uterine, vaginal, nasal and pulmonary microbiota from 16 captive hamadryas baboons by culture-independent sequencing of the 16S rRNA gene hypervariable V3-V4 region. Our study gives detailed insights into the baboon microbiome structure and ecology.

## Materials and methods

### Controls manage the risk of contamination during wet-lab processing and sterile surgical procedure manage the risk of contamination at sampling

To avoid contamination risks, we strictly controlled the sampling process, DNA extraction, PCR and sequencing. All DNA extractions strictly followed the aseptic operation process under a clean bench. We also have DNA extraction negative control (from the DNA extraction to the sequencing process), sequencing blank control (clean water for sequencing), and sequencing positive control (mock community, *E.coli*). The detailed information is included in the supplementary material.

### Sample collection

Samples were collected from 16 captive baboons (*Papio hamadryas*) housed at the Copenhagen Zoo, Denmark. Eight different sites were sampled (Fig.1) and all sample information is listed in Table S4. Animals were anesthetized for a full medical evaluation and physical examination. Non-invasive samples (vagina, nose, oral, oropharynx, and gut) were collected in a sheltered housing facility using sterile polyester swabs (cat no. 300263, Deltalab, Spain). Following thorough medical evaluations, eight of the animals were euthanized by a licensed veterinarian and a thorough postmortem examination was conducted in a separate necropsy room. Carcasses were opened ventrally to expose the organs, and all invasive sampling was performed sequentially from cranial to caudal. Lungs were excised, and both the left and right main bronchi were swabbed, whereas for all other invasive samples, a small surgical incision was made to obtain the swabs. A new sterile scalpel was used for each organ; new gloves were donned and new surgical utensils were used for each of the carcasses. Permission to sample and euthanize the animals was granted by the Copenhagen Zoo.

**Fig 1.**
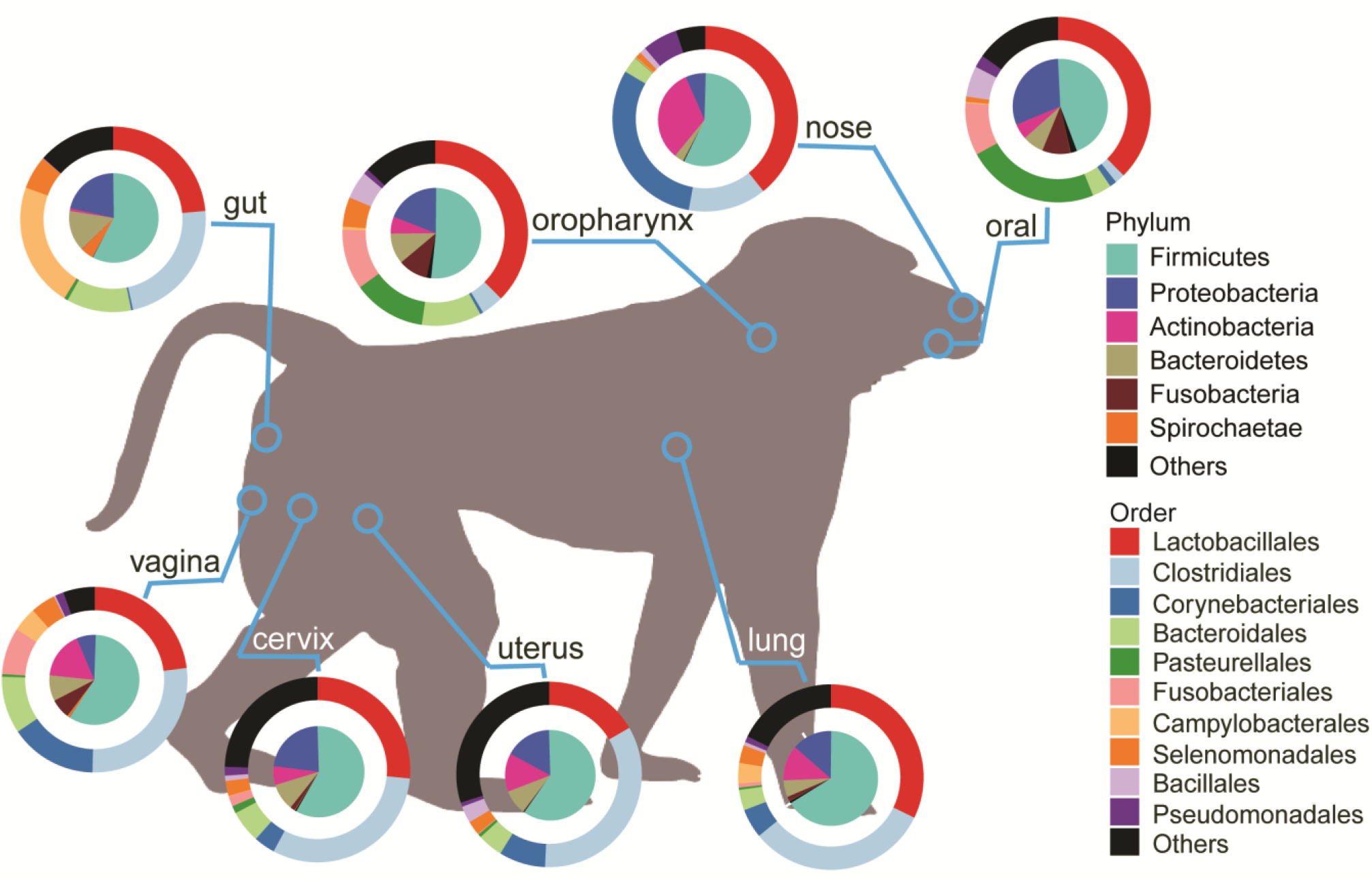
Microbial composition on phylum and order levels of each anatomic site. Pie charts show the average microbial composition of eight body habitats on order (pie chart) and phylum level (donut chart), respectively. *Lactobacillales* was found in all the body sites (Mean 32% ± SD 15%). The microbial composition for each individual baboon can be found in Supplementary Fig.S1.

### DNA extraction and 16S rRNA gene amplicon sequencing

Genomic DNA was extracted from swab samples with the PowerLyzer® PowerSoil® DNA Isolation Kit (MO-BIO Laboratories, Inc., Carlsberg, CA, USA), and 50 μL of elution buffer was used for each sample. All operations were performed under aseptic conditions. Extracted DNA was stored at −20°C. Sterilized PBS solution and Molecular grade water (Sigma-Aldrich, United States) were used as DNA extraction and DNA amplification negative control, whereas mock community and only E.coli strain were included in all the following steps as a positive control. The 16S rRNA gene hypervariable V3-V4 region was amplified with 2 μl template DNA, using 0.25 μl Phusion high-fidelity (HF) DNA Polymerase (Thermo Fisher Scientific, Waltham, MA, USA), 5 μl 5×Phusion buffer HF, 0.5 μl 10mM dNTPs, 1 μl 10 μM of each primer (the modified broad primers 341F (5’-CCTAYGGGRBGCASCAG-3’) and Uni806R (5’-GGACTACNNGGGTATCTAAT-3’) [40] in a 25 μl PCR reaction volume. The first PCR program included 30 s at 98°C, 30 cycles of 5 s at 98°C, 15 s at 56°C, and 72°C for 10 s, and then 5 min at 72°C. In the second PCR, sequencing primers and adaptors were attached to the amplicon library following the first PCR conditions with only 15 cycles. The size of the PCR product (≈466 bp) was evaluated using gel electrophoresis. The amplicon products were purified by use of Agencourt AMPure XP beads (Beckman Coulter Genomics, MA, USA) with the 96-well magnet stand, normalized by the SequalPrepTM Normalization Plate (96) kit (Invitrogen Ltd., Paisley, UK), pooled in equimolar concentrations and concentrated using the DNA Clean & Concentrator™-5 Kit (Zymo Research, Irvine, CA, USA). Sequencing of the amplicon library was performed on the Illumina MiSeq System with MiSeq reagent kit v2 (Illumina Inc., CA, USA), including 5.0% PhiX as an internal control.

### Bioinformatics analysis and Statistical analysis

The raw fastq files were demultiplexed using the Miseq Controller Software. Primers and diversity spacers were removed from fastq files using “Cutadapt” [41]. The data trimming and feature classification were done using Qiime 2 Core 2017.12 distribution microbiota analysis platform (https://qiime2.org). Paired-end sequences were merged by vsearch plugin [42] and then followed by filtering with the quality-filter plugin [43], both with default settings. Deblur plugin was then used to denoise the sequences with a trim length of 400bp based on quality score plots [44]. Sequence alignments were generated using MAFFT and the aligned sequences were masked by MASK plugin [45]. FastTree and midpoint-root built-in phylogeny plugin were used to create a rooted phylogenetic tree [46]. Pre-fitted sklearn-based taxonomy classifier (https://github.com/qiime2/q2-feature-classifier) was used to blast representative sequences against silva 132 database for taxonomic classification of features [47]. Rarefaction curve was plotted by alpha_rarefaction.py workflow in Qiime 2.

The distribution histogram of average unweighted UniFrac distances between each sample and all the rest samples were plotted to confirm that PCR and Sequencing controls differed from our samples (supplementary material). The DNA amplification negative control with only 63 reads was filtered. The histogram showed three controls (DNA extraction blank control, *E.coli* strain and Mock control) were far away from the real samples, which meant controls are different from the real samples. The rarefaction curves (supplementary material) demonstrated the observed richness in a given count of sequences. Observed richness curves reached asymptotes after 4000 reads for most samples. With an average of 15,247 clean sequences per sample, sufficient sequences for all 184 samples were generated to characterize the microbial community in the eight body habitats.

The open-source statistical program “R” was used for data treatment and statistical analysis [48], predominantly the *R*-package “phyloseq” [49]. Alpha diversity between the groups was tested by analysis of variance using the function “anova”. If significant differences between the groups were present, multiple comparisons with the function “TukeyHSD” were performed pairwise between all groups (all three functions from *R*-package “stats”). Bray Curtis distance was used to explain differences among microbial communities and the dissimilarity was examined by permutational multivariate analysis of variance (PERMANOVA, “vegan” function “adonis”) [50]. *R* Function “pairwise.adonis” [51] was used for multiple comparisons and Bonferroni correction was used to account for multiple comparisons. Group Divergence was quantified as the average dissimilarity of each sample from the group mean by using the function “divergence” from R-package “microbiome” [52]. Venn diagram was plotted by *R* function “VennCounts” and “VennDiagram” in *R* package “VennDiagram” [53]. *R* function “rcorr” was used to compute the Spearman correlation analysis and the significance levels [54]. The correlation matrix and the significance test were visualized by *R* function “corrplot” [55]. Pie chart, donut chart, violin plot, box plot, heatmap, bar chart and circle bar chart were plotted using ggplot2 [56].

## Results

### Microbial distribution in different body sites of captive baboons

Six main phyla were detected in the eight body sites; of them, the phylum *Spirochaetae* was found to be abundant in the baboon gut (Fig.1). *Firmicutes* dominated in all body sites. The dominance of other phyla varied in the body sites; for instance, *Fusobacteria* was dominant in the oral cavity and oropharynx, and *Actinobacteria* in the nose. On the order level, we found *Lactobacillales* to be abundant in all eight body sites (Mean 32% ± SD 15%), especially in the oral cavity, oropharynx and nose, constituting around 40% of the total bacteria. In addition to *Lactobacillales*, *Clostridiales* were predominant in the vagina, cervix, uterus, lungs, and gut, together accounting for 47-58% of the microbiota. The oral and oropharynx shared a microbial profile mainly composed of *Lactobacillales*, *Bacteroidales*, and *Pasteurellales*, representing over 60% of the microbiota. *Lactobacillales* and *Corynebacteriales* were the two major bacteria in the nose, together comprising almost 70% of the microbiota.

### Microbial characterizations varied in the body sites

To investigate microbial features of different body sites, we analyzed microbial diversity between and within the different body sites, quantified the bacterial divergence within each body site and compared gut microbial diversity between and within maternal lines (Fig.2). Non-metric multidimensional scaling (NMDS) based on a Bray-Curtis distance matrix (Fig.2A) showed that the oral cavity, oropharynx, and nose had unique microbial profiles but the microbial profiles in the cervix, uterus, vagina, rectum, and lungs did not have significant differences (pairwise comparison, adjusted *p* values were listed in Table S1, Permutational Multivariate Analysis of Variance). Microbial diversity in the baboon nose, oral cavity and oropharynx was significantly lower than in the lungs and reproductive tract (adjusted *p* values were listed in Table S2, TukeyHSD) (Fig.2B). Divergence of microbes in a specific body site was quantified and extrapolated in Fig.2C. The microbes in baboon nose and oropharynx had a smaller spread but were more heterogeneous in the reproductive tract, oral and lungs (adjusted *p* values were listed in Table S3, Pairwise Wilcoxon Rank Sum Tests). The vertical inheritance of gut microbial communities was analyzed on the three maternal lines in the present cohort by checking whether hosts of the same maternal line shared more bacterial phylotypes on average than unrelated hosts (Fig.2D). Statistical analysis demonstrated that the microbial phylotypes on the maternal lines were similar to those of unrelated individuals (Wilcoxon test, *p* = 0.776), indicating the vital role of horizontal exchange in shaping the gut microbiota.

**Fig 2.**
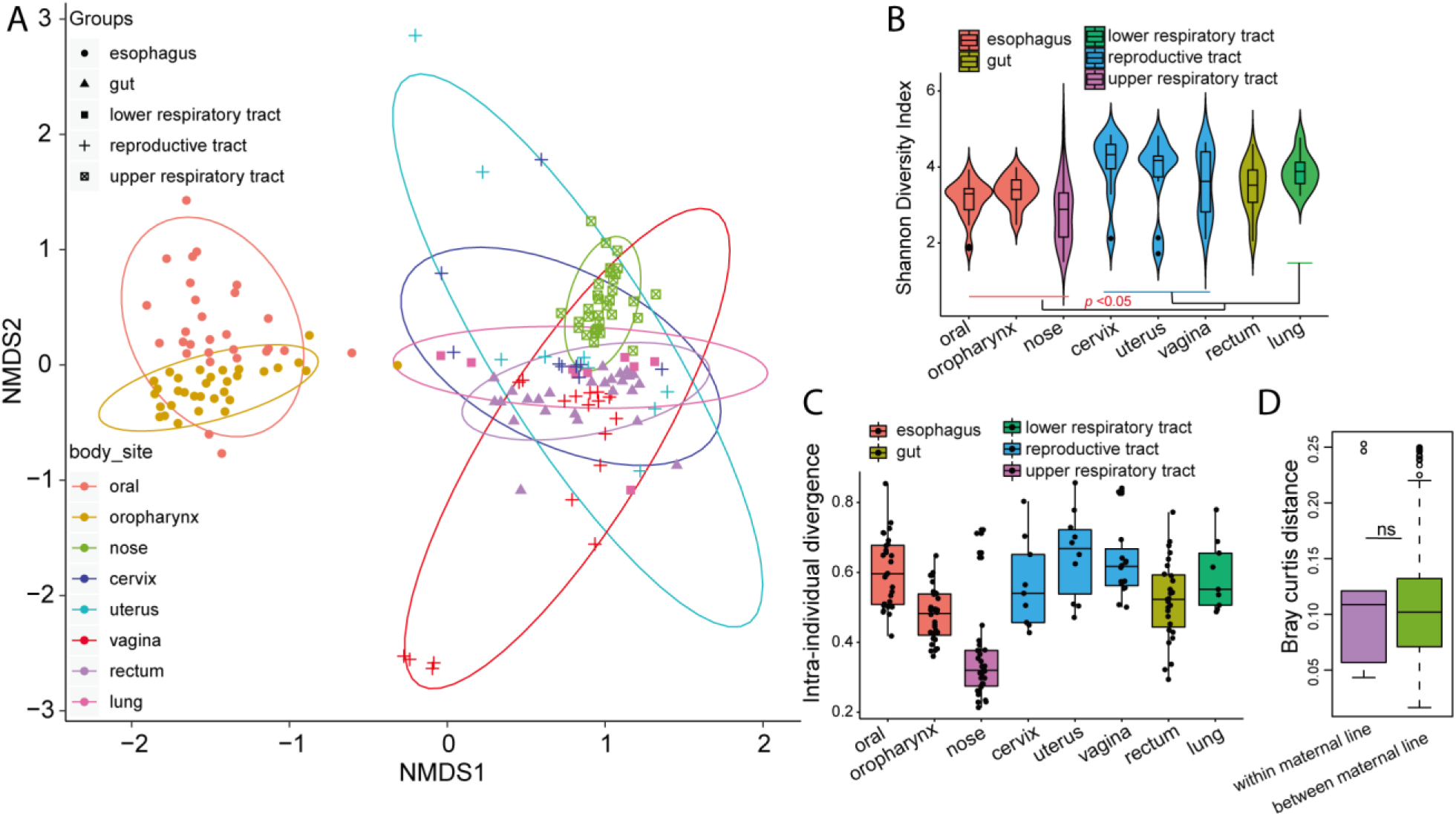
Microbial characterization in different body sites. A) Non-metric multidimensional scaling (NMDS) based on Bray-Curtis distance of microbial communities from the baboon oral cavity, oropharynx, nose, cervix, uterus, vagina and rectum. The colored lines surrounding each sample type are covariance ellipsoids. B) Alpha diversity in different body habitats, grouped by area, as measured using the Shannon index of ASV-level bacteria. Pharynx and nose had a significantly lower diverse microbiota than lungs and reproductive tract (TukeyHSD). C) Divergence of microbes in a specific body site was quantified as the average dissimilarity of each sample from the group mean. D) Bray curtis distance within and between maternal lines for gut microbiota. ns means no significant difference detected between the same maternal line and different maternal lines by Wilcoxon test (*p*=0.776).

### Baboons had significant different vagina, gut and oral microbiomes from human

The 1041 comparable Miseq sequencing data from the human oral cavity, gut and vagina [17] were reanalyzed using the same pipeline in this study. We found that the baboon microbiomes in the three body sites were significantly different from human microbiomes (pairwise comparison, *p* = 0.015* for oral vs oral microbiome, *p* = 0.015* for gut vs gut microbiome, *p* = 0.015* for vaginal vs vaginal microbiome, Permutational Multivariate Analysis of Variance, Fig.3A). Besides, the vaginal microbiome of baboon was more diverse than that of the human (adjusted *p*-value < 1e−7***, TukeyHSD, Fig.3B). The human had a slightly higher diversity of oral microbiome than baboon and a similar diversity of gut microbiome to the baboon. *Lactobacillus* (Mean 13% ± SD 12%) was not dominant in the baboon vagina as in the human vagina (Mean 54% ± 37%), but it was the most abundant genus in the baboon gut (Mean 22% ± SD 17%) (Fig.3C). By contrast, the human gut only contained an average relative abundance of 0.35% *Lactobacillus* (Fig.3C). In spite of an overall significant difference in oral microbial profiles between baboon and human (Fig.3A), the 30 most abundant genera in their oral cavity were similar in the sample dendrogram (Fig.3C). *Streptococcus* was the most abundant genus both in the human and baboon oral cavity.

**Fig 3.**
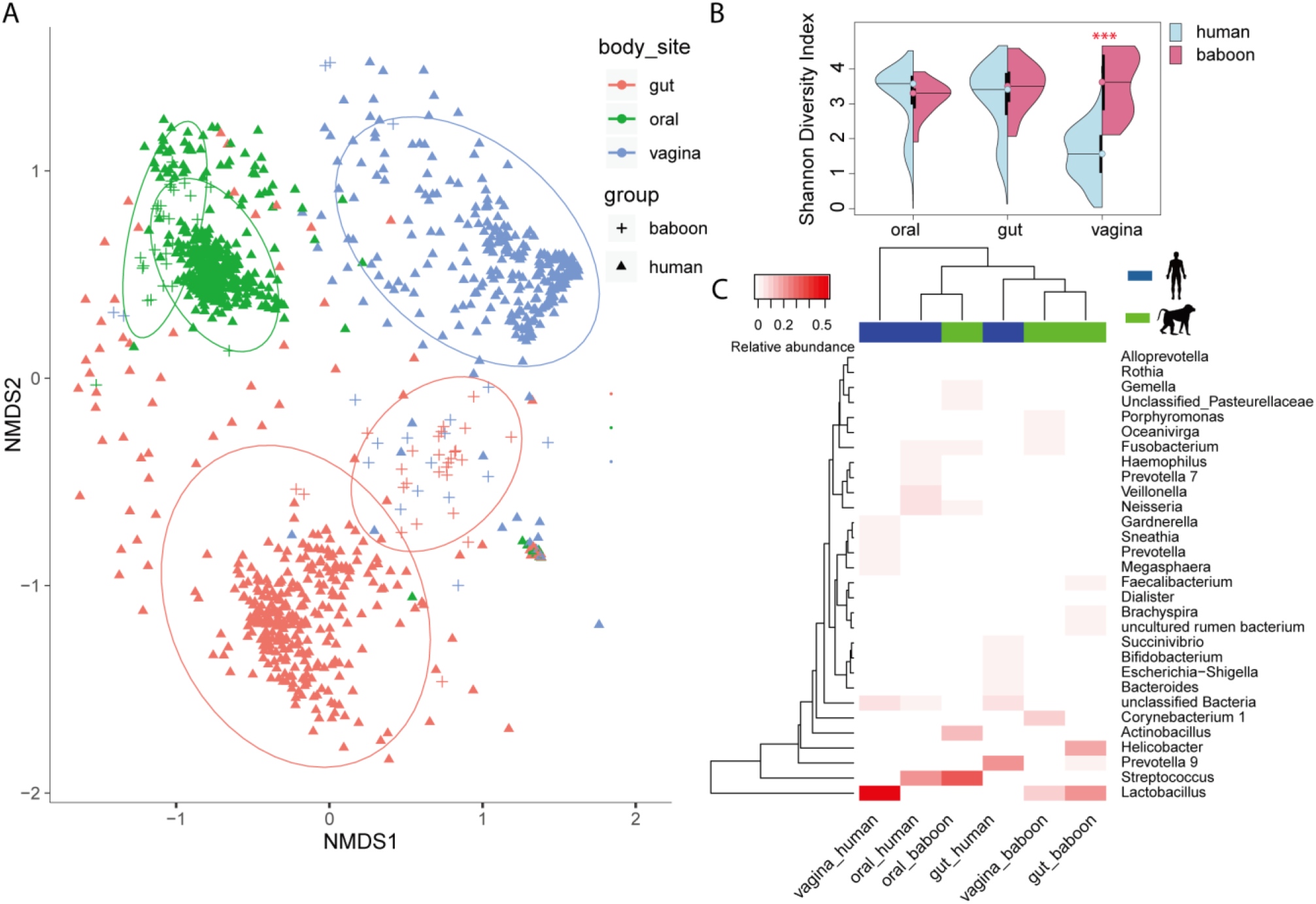
Comparisons between the human and baboon microbiomes in the oral cavity, gut and vagina. A) Non-metric multidimensional scaling (NMDS) based on a Bray-Curtis distance matrix of microbial communities from baboon and human oral, gut and vagina. The colored lines surrounding each sample type are covariance ellipsoids. B) Microbial alpha diversity in the human and baboon gut, oral cavity and vagina, as measured using the Shannon index of ASV-level bacteria. C) Heatmap showing the 30 most abundant bacterial genera in human and baboon oral, nose and vagina.

### The microbes of group-living baboons shared more similarities across body sites than human

As we know, bacterial exchange during group living is inevitable. We define those ASVs which are present in at least one baboon sample in each body site, as shared ASVs. The different body sites of group-living baboons shared more ASVs than those of humans (Fig.4A). These shared ASVs did not show any significant correlations in humans but had a significant positive correlation of gut and vagina in the baboon (Fig.4A), indicating the potential bacterial exchange. Of the 15 shared ASVs by the human oral cavity, gut and vagina, 5 ASVs belonged to unclassified taxa (Fig.4A). Most shared ASVs by the baboon’s oral cavity, gut and vagina belonged to *Firmicutes* (Fig.4A). In addition, the human oral cavity, gut and vagina had its own unique microbial compositions as shown in Non-metric multidimensional scaling (NMDS) plot (pairwise comparison, *p* = 0.003** for oral vs gut microbiome, *p* = 0.003** for oral vs vaginal microbiome, *p* = 0.003** for gut vs vaginal microbiome, Permutational Multivariate Analysis of Variance, Fig.4A). However, the baboon gut microbiome was similar to the vaginal microbiome (Fig.2A) in spite of a big variance within an individual (Fig.4B). From Figure 2A, it is clear that multiple body sites of baboons had similar microbial profiles. Considering the potential bacterial exchange across body sites among co-habitants, we analyzed the shared ASVs (Fig.4C) and their correlations (Fig.4D) of all the 8 body sites. These shared 33 ASVs contributed to 32% (from 17-70% in each body habitat) of the mean relative abundance (Fig.4C) and *Firmicutes* accounted for the most shared phylum (Fig.3A). Among these shared 33 ASVs, 11 ASVs belong to the genus *Lactobacillus* and 2 ASVs belong to the genus *Faecalibacterium*. In addition, apart from the oral, oropharynx and nose sites, these shared 33 ASVs were positively correlated with each other.

**Fig 4.**
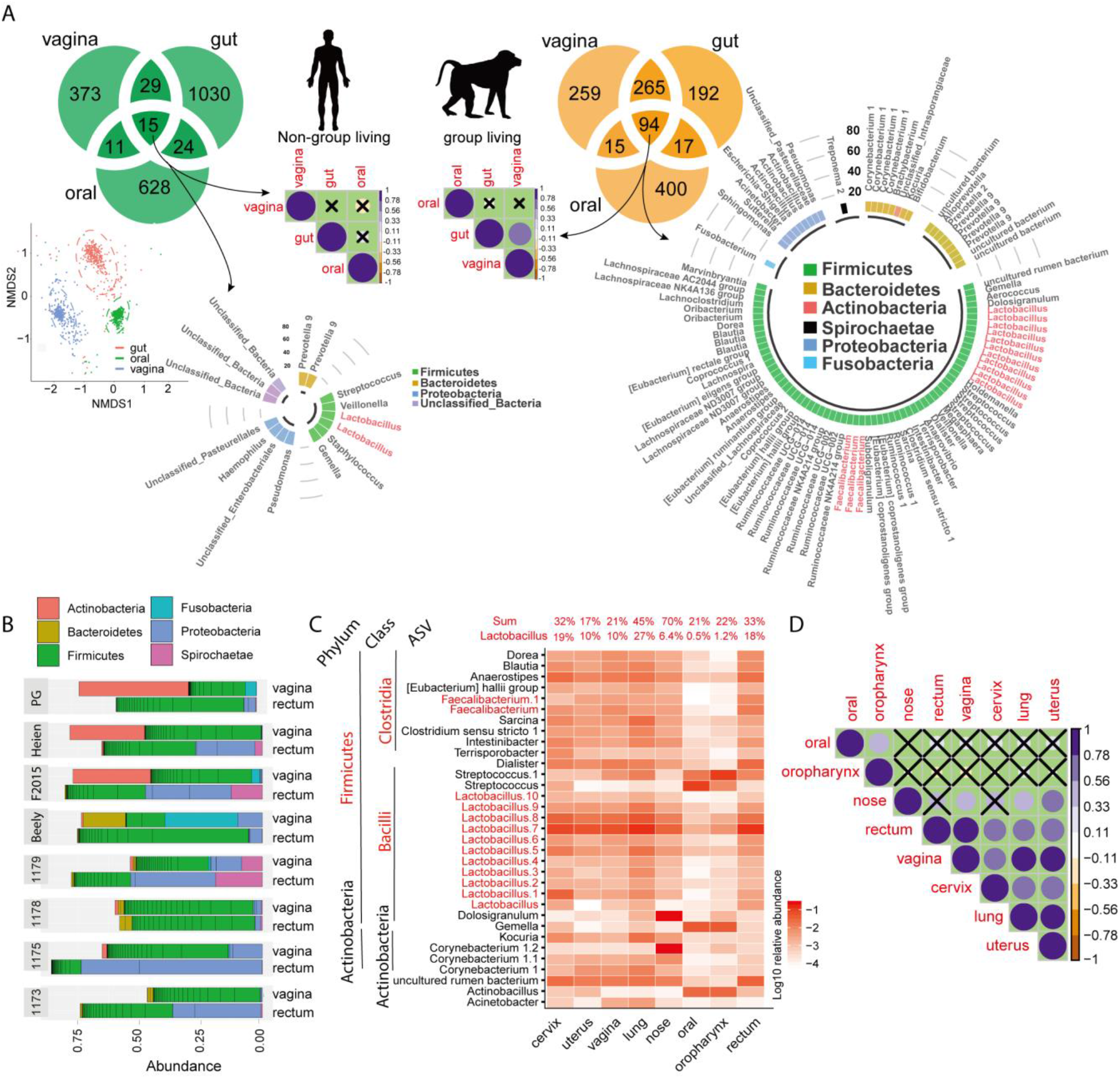
Shared microbial ASVs and their correlation analysis among human/baboon oral, gut and vagina, and among all eight body sites of baboons. A) Venn diagrams showing the number of shared ASVs and spearman correlation matrix for shared ASVs among oral, gut and vagina of humans and baboons. X in the correlation matrix means no significant correlations. All the significant correlations shown in the matrix have an adjusted *p*-value<0.01. Color intensity and the size of the circle are proportional to the correlation coefficients. The shared ASVs by human and baboon samples were plotted in the circular barplots. Bray-Curtis distance of microbial communities from human oral, gut and vagina was shown in NMDS plot. B) Bar plots showing the 40 most abundant bacteria ASVs of vagina and rectum in eight baboons. C) Heatmap for 33 ASVs shared by all body sites. The ASVs belonging to genus *Lactobacillus* and *Faecalibacterium* are marked in red. The relative abundances of *Lactobacillus* and the 33 ASVs are listed. D) Spearman correlation analysis for 33 ASVs shared by eight body sites. Only the Spearman correlation coefficients (PCC) with an adjusted *p*<0.01 were plotted. X in the correlation matrix means no significant correlations. Color intensity and the size of the circle are proportional to the correlation coefficients.

## Discussion

In our study, we find that the baboon microbiome has unique characteristics compared to the human microbiome. Group-living baboons shared more similarities among body sites than humans. A highly positive correlation of those shared ASVs among the different body sites suggested the potential bacterial exchange throughout the body. Captive baboons investigated in this study live in the same environment and shared the same diet from birth to death. Generally, group-living entails frequent social interactions, especially for highly social baboons [18]. Frequent social interactions (mostly grooming) are necessary for baboons to maintain affiliative bonds [11]. Bacterial exchange in a shared household has frequently been reported among primates. Members of a shared household had a more similar gut and skin bacterial communities than individuals living in separate households, indicating that a shared lifestyle or environment affected the microbiota composition [19, 20]. Similarly, in a study of the gut microbiota in wild chimpanzees, group-living baboons shared more of their gut microbiome than individuals from different groups [21]. In the study of baboon gut microbiota by Tung et al., baboon gut microbiome was detected to have a high group specificity as well [22]. They found even if two social groups of wild baboons shared almost the same diet, the gut microbiota of these two groups were still significantly different and shaped by the social interactions [22]. Therefore, we speculate that bacterial exchange was an important reason for microbial similarities in the multiple body sites of group-living baboons. In addition, the gut microbiota within the three maternal lines did not show a significant difference in comparison to those of unrelated individuals, which indicated that the gut microbiota is more affected by bacterial exchange. Our findings were consistent with the results previously reported in the literature [21]: inheritance of microbial communities across generations were primarily horizontal among interacting hosts. In the present study, approximately one-third of all the shared OTUs were from the genus *Lactobacillus* and *Faecalibacterium*, which tended to be shared between body sites. Many species from *Lactobacillus* and *Faecalibacterium* are widely considered to be probiotics [23, 24] which could potentially help explain the driving force for bacterial transmission.

The polygynous mating system and promiscuity of the baboons boosted the probability of genital bacterial transfer. In this study, we found that the baboon vaginal microbiome was similar to the gut microbiome, which is different from human counterparts. For baboons, some grooming bouts, especially those directed from adult males to estrous females, concentrate heavily on the anogenital region, increasing the probability of fecal-vagina transfer [25]. Besides, we found that the baboon vagina had a distinct microbiota profile from that of humans (Fig.3A). Baboon vagina had a relatively high microbial diversity (Fig.3B) and *Lactobacillus* was not so predominated in baboon vagina as in humans (Fig.3C). Similar findings have been reported in other studies [13]. The human vagina is primarily colonized by *Lactobacillus* [26], which maintains an acidic environment and prevents the invasion of nonindigenous strains and potential pathogens and could even account for 65.9% to 98.1% of the vaginal microbiota [27–30]. However, to some extent, the baboon vaginal microbial profiles characterized by low *Lactobacillus* abundance, low lactic acid concentration and a higher, near-neutral vaginal pH [31], typically associated with bacterial vaginosis in the human vagina [32].

Compared to the gut microbiota, the lung microbiota of nonhuman primates is still poorly understood. In the present study, the lung samples shared a similar bacterial profile with the genital region (Fig.2A), which was in agreement with other studies using murine studies [16]. Barfod *et al*. proposed that the core lung microbiota of mice is established in utero, during and after birth in the very early life, similar to the gut microbiota in humans [16]. In another research, maternal inheritance has been shown to play a vital role in the bacterial community composition of the lungs in early life [33], whereas the lung microbiota of adults demonstrated higher resilience towards environmental variations [34]. Therefore, the same living environment could be the reason for the microbial similarities between the lung and reproductive tract. Studies of the lung microbiome suffer from low amounts of available DNA, and this has been shown to lead to contamination and inflated diversity estimates [35]. This study also saw low concentrations, so to prevent these problems in the current work we have done a detailed analysis of negative sequencing controls (supplementary material). Additionally, we have detailed descriptions of the surgical procedures and steps taken to avoid contamination by this route.

The baboon oral cavity had a unique microbial distribution and exhibited minor overlap with other body habitats (Fig.2A); it had somewhat lower microbial diversity than humans (Fig.3B), in spite of the fact that humans exhibit more oral hygiene practices. This could be because the baboon diet in captivity was still fairly simpler compared to humans. The baboon gut microbiome is unique compared to the human gut microbiome. *Spirochaetae*, which was extremely rare in the modern human but enriched in ancient humans [36], was also enriched in the baboon gut (Fig.1). *Lactobacillus* was the most abundant genus in the baboon gut but presented a low abundance in the human gut (Fig.3C). Notably, the important human gut bacterium, *Akkermansia* [37], was not detected in the baboon gut. Therefore, *Akkermansia* was more like human-specific and thus absent in the baboons in our study, no *Akkermansia* has been reported in NHPs [38, 39].

## Conclusion

To our knowledge, this study is the first to test microbial compositions in cohabitating baboons across different body sites. In summary, our results showed that baboons have a unique microbiome compared to humans. The microbial diversity of the baboon vagina was much higher than that of humans. *Lactobacillus* was not so predominant in baboon vagina as in the human vagina but was the most abundant genus in the baboon gut. The microbial compositions in the baboon reproductive tract, gut and lungs were similar. Oral cavity, vagina and gut in group-living baboons shared more bacterial ASVs than humans. The significantly positive correlations of those shared ASVs between multiple body sites in this group of baboons combined with highly social characteristics of baboons indicated a potential bacterial exchange throughout the body. We reported that the probably transmitted bacteria across body sites tend to be bacteria known to be beneficial in humans, which may suggest that some modern human populations, due to changed social behaviors, may have lost an important source of beneficial microbiota with consequences for human health.

## Acknowledgments

This work was supported by the China Scholarship Council and by the Danish Council for Independent Research grant. The authors declare that they have no competing interests. We express our deepest gratitude to Copenhagen Zoo for collaboration with the baboon samples. We acknowledge Luma Odish for the help and support with the construction of the 16S rRNA gene amplicon libraries and sequencing.

## Data availability

The dataset analyzed during the current study are available in the Sequence Read Archive (SRA) repository, http://www.ncbi.nlm.nih.gov/bioproject/464237.

---

**Fig S1.** Bar plot shows the microbial composition of eight body habitats on order level. *Lactobacillales* were present in all the body sites.

**Table S1**: Sample overview.

**Supplementary material**: controls manage risk of contamination during wet-lab processing and sterile surgical procedure manage risk of contamination at sampling.

## References

1. Cho I, Blaser MJ. The human microbiome: At the interface of health and disease. Nat Rev Genet 2012; 13: 260–270.

2. Pflughoeft KJ, Versalovic J. Human Microbiome in Health and Disease. Annu Rev Pathol Mech Dis 2012; 7: 99–122.

3. Ezenwa VO, Gerardo NM, Inouye DW, Medina M, Xavier JB. Animal behavior and the microbiome. Science(80-). 2012., 338: 198–199

4. Nih T, Working HMP. The NIH Human Microbiome Project. Genome Res 2009; 19: 2317–2323.

5. Tung J, Barreiro LB, Burns MB, Grenier JC, Lynch J, Grieneisen LE, et al. Social networks predict gut microbiome composition in wild baboons. Elife 2015; 2015.

6. Moeller AH, Foerster S, Wilson ML, Pusey AE, Hahn BH, Ochman H. Social behavior shapes the chimpanzee pan-microbiome. Sci Adv 2016; 2: e1500997.

7. Rogers J, Hixson JE. Baboons as an animal model for genetic studies of common human disease. Am J Hum Genet 1997; 61: 489–493.

8. Barrett L, Henzi SP. Baboons. Curr Biol 2008; 18: 404–406.

9. Swedell L, Plummer T. A Papionin Multilevel Society as a Model for Hominin Social Evolution. Int J Primatol 2012; 33: 1165–1193.

10. Newman TK, Jolly CJ, Rogers J. Mitochondrial phylogeny and systematics of baboons (Papio). Am J Phys Anthropol 2004; 124: 17–27.

11. Lehmann J, Korstjens AH, Dunbar RIM. Group size, grooming and social cohesion in primates. Anim Behav 2007; 74: 1617–1629.

12. Mckenney EA, Ashwell M, Lambert JE, Fellner V. Fecal microbial diversity and putative function in captive western lowland gorillas (Gorilla gorilla gorilla), common chimpanzees (Pan troglodytes), Hamadryas baboons (Papio hamadryas) and binturongs (Arctictis binturong). Integr Zool 2014; 9: 557–569.

13. Yildirim S, Yeoman CJ, Janga SC, Thomas SM, Ho M, Leigh SR, et al. Primate vaginal microbiomes exhibit species specificity without universal Lactobacillus dominance. ISME J 2014; 8: 2431–2444.

14. Miller EA, Livermore JA, Alberts SC, Tung J, Archie EA. Ovarian cycling and reproductive state shape the vaginal microbiota in wild baboons. Microbiome 2017; 5: 1–14.

15. Beck JM, Young VB, Huffnagle GB. The microbiome of the lung. Transl Res 2012; 160: 258–266.

16. Barfod KK, Roggenbuck M, Hansen LH, Schjørring S, Larsen ST, Sørensen SJ, et al. The murine lung microbiome in relation to the intestinal and vaginal bacterial communities. BMC Microbiol 2013; 13.

17. Bisanz JE, Enos MK, PrayGod G, Seney S, Macklaim JM, Chilton S, et al. Microbiota at multiple body sites during pregnancy in a rural tanzanian population and effects of Moringa-supplemented probiotic yogurt. Appl Environ Microbiol 2015; 81: 4965–4975.

18. Schreier AL, Swedell L. The fourth level of social structure in a multi-level society: Ecological and social functions of clans in Hamadryas Baboons. Am J Primatol 2009.

19. Rothschild D, Weissbrod O, Barkan E, Kurilshikov A, Korem T, Zeevi D, et al. Environment dominates over host genetics in shaping human gut microbiota. Nat Publ Gr 2018; 555: 210–215.

20. Song SJ, Lauber C, Costello EK, Lozupone CA, Humphrey G, Berg-Lyons D, et al. Cohabiting family members share microbiota with one another and with their dogs. Elife 2013; 2013.

21. Moeller AH, Foerster S, Wilson ML, Pusey AE, Hahn BH, Ochman H. Social behavior shapes the chimpanzee pan-microbiome. Sci Adv 2016; 2.

22. Tung J, Barreiro LB, Burns MB, Grenier JC, Lynch J, Grieneisen LE, et al. Social networks predict gut microbiome composition in wild baboons. Elife 2015; 2015: 2013–2015.

23. Liévin-Le Moal V, Servin AL. Anti-infective activities of Lactobacillus strains in the human intestinal microbiota: From probiotics to gastrointestinal anti-infectious biotherapeutic agents. Clin Microbiol Rev 2014; 27: 167–199.

24. Miquel S, Martín R, Rossi O, Bermúdez-Humarán LG, Chatel JM, Sokol H, et al. Faecalibacterium prausnitzii and human intestinal health. Curr Opin Microbiol 2013; 16: 255–261.

25. Tung J, Barreiro LB, Burns MB, Grenier JC, Lynch J, Grieneisen LE, et al. Social networks predict gut microbiome composition in wild baboons. Elife 2015; 2015: 1–18.

26. Srinivasan S, Fredricks DN. The Human Vaginal Bacterial Biota and Bacterial Vaginosis. Interdiscip Perspect Infect Dis 2008; 2008: 1–22.

27. Kim TK, Thomas SM, Ho M, Sharma S, Reich CI, Frank JA, et al. Heterogeneity of vaginal microbial communities within individuals. J Clin Microbiol 2009; 47: 1181–1189.

28. Hummelen R, Fernandes AD, Macklaim JM, Dickson RJ, Changalucha J, Gloor GB, et al. Deep sequencing of the vaginal microbiota of women with HIV. PLoS One 2010; 5.

29. Srinivasan S, Liu C, Mitchell CM, Fiedler TL, Thomas KK, Agnew KJ, et al. Temporal variability of human vaginal bacteria and relationship with bacterial vaginosis. PLoS One 2010; 5.

30. Ravel J, Brotman RM, Gajer P, Ma B, Nandy M, Fadrosh DW, et al. Daily temporal dynamics of vaginal microbiota before, during and after episodes of bacterial vaginosis. Microbiome 2013; 1: 29.

31. Miller EA, Beasley DAE, Dunn RR, Archie EA. Lactobacilli dominance and vaginal pH: Why is the human vaginal microbiome unique? Front Microbiol 2016; 7: 1–13.

32. Aldunate M, Srbinovski D, Hearps AC, Latham CF, Ramsland PA, Gugasyan R, et al. Antimicrobial and immune modulatory effects of lactic acid and short chain fatty acids produced by vaginal microbiota associated with eubiosis and bacterial vaginosis. Front Physiol. 2015., 6

33. Mortensen MS, Brejnrod AD, Roggenbuck M, Abu Al-Soud W, Balle C, Krogfelt KA, et al. The developing hypopharyngeal microbiota in early life. Microbiome 2016; 4: 70.

34. Kostric M, Milger K, Krauss-Etschmann S, Engel M, Vestergaard G, Schloter M, et al. Development of a Stable Lung Microbiome in Healthy Neonatal Mice. Microb Ecol 2018; 75: 529–542.

35. Salter SJ, Cox MJ, Turek EM, Calus ST, Cookson WO, Moffatt MF, et al. Reagent and laboratory contamination can critically impact sequence-based microbiome analyses. BMC Biol 2014; 12: 1–12.

36. Turroni S, Rampelli S, Centanni M, Schnorr SL, Consolandi C, Severgnini M, et al. Enterocyte-associated microbiome of the Hadza hunter-gatherers. Front Microbiol 2016; 7: 1–12.

37. van Passel MWJ, Kant R, Zoetendal EG, Plugge CM, Derrien M, Malfatti SA, et al. The genome of Akkermansia muciniphila, a dedicated intestinal mucin degrader, and its use in exploring intestinal metagenomes. PLoS One 2011; 6.

38. Li X, Liang S, Xia Z, Qu J, Liu H, Liu C, et al. Establishment of a Macaca fascicularis gut microbiome gene catalog and comparison with the human, pig, and mouse gut microbiomes. Gigascience 2018; 7: 1–10.

39. Firrman J, Liu LS, Tanes C, Bittinger K, Mahalak K, Rinaldi W. Metagenomic assessment of the Cebus apella gut microbiota. Am J Primatol 2019; 1–11.

40. Takai K, Horikoshi K, Takai KEN. Rapid Detection and Quantification of Members of the Archaeal Community by Quantitative PCR Using Fluorogenic Probes Rapid Detection and Quantification of Members of the Archaeal Community by Quantitative PCR Using Fluorogenic Probes. Appl Environ Microbiol 2000; 66: 5066–5072.

41. Martin M. Cutadapt removes adapter sequences from high-throughput sequencing reads. EMBnet.journal 2011; 17: 10.

42. Rognes T, Flouri T, Nichols B, Quince C, Mahé F. VSEARCH: a versatile open source tool for metagenomics. PeerJ 2016; 4: e2584.

43. Bokulich NA, Subramanian S, Faith JJ, Gevers D, Gordon JI, Knight R, et al. Quality-filtering vastly improves diversity estimates from Illumina amplicon sequencing. Nat Methods 2013; 10: 57–59.

44. Amir A, McDonald D, Navas-Molina JA, Kopylova E, Morton JT, Zech Xu Z, et al. Deblur Rapidly Resolves Single-Nucleotide Community Sequence Patterns. mSystems 2017; 2: e00191–16.

45. Katoh K, Standley DM. MAFFT multiple sequence alignment software version 7: Improvements in performance and usability. Mol Biol Evol 2013; 30: 772–780.

46. Price MN, Dehal PS, Arkin AP. FastTree 2 - Approximately maximum-likelihood trees for large alignments. PLoS One 2010; 5.

47. Quast C, Pruesse E, Yilmaz P, Gerken J, Schweer T, Yarza P, et al. The SILVA ribosomal RNA gene database project: Improved data processing and web-based tools. Nucleic Acids Res 2013; 41.

48. Hornik K. The Comprehensive R Archive Network. Wiley Interdiscip Rev Comput Stat. 2012., 4: 394–398

49. McMurdie PJ, Holmes S. Phyloseq: An R Package for Reproducible Interactive Analysis and Graphics of Microbiome Census Data. PLoS One 2013; 8.

50. Oksanen J, Kindt R, Legendre P, O’Hara B, Simpson GL, Solymos PM, et al. The vegan package. Community Ecol Packag 2008; 190.

51. Arbizu M. Pairwiseadonis: Pairwise multilevel comparison using adonis. R Packag Version 00 2017; 1.

52. Leo Lahti, Sudarshan Shetty et al. (2017). Tools for microbiome analysis in R. Version 1.9.19. URL: http://microbiome.github.com/microbiome

53. Chen H, Boutros PC. VennDiagram: A package for the generation of highly-customizable Venn and Euler diagrams in R. BMC Bioinformatics 2011; 12.

54. Miscellaneous TH, Yes L. Package ‘Hmisc’. 2018.

55. Wei T, Simko V. The corrplot package. R Core Team 2016.

56. Wilkinson L. ggplot2: Elegant Graphics for Data Analysis by WICKHAM, H. Biometrics 2011; 67: 678–679.

