## Supplementary figures and images for "The microbiome of captive hamadryas baboon"

### Supplementary Fig.1

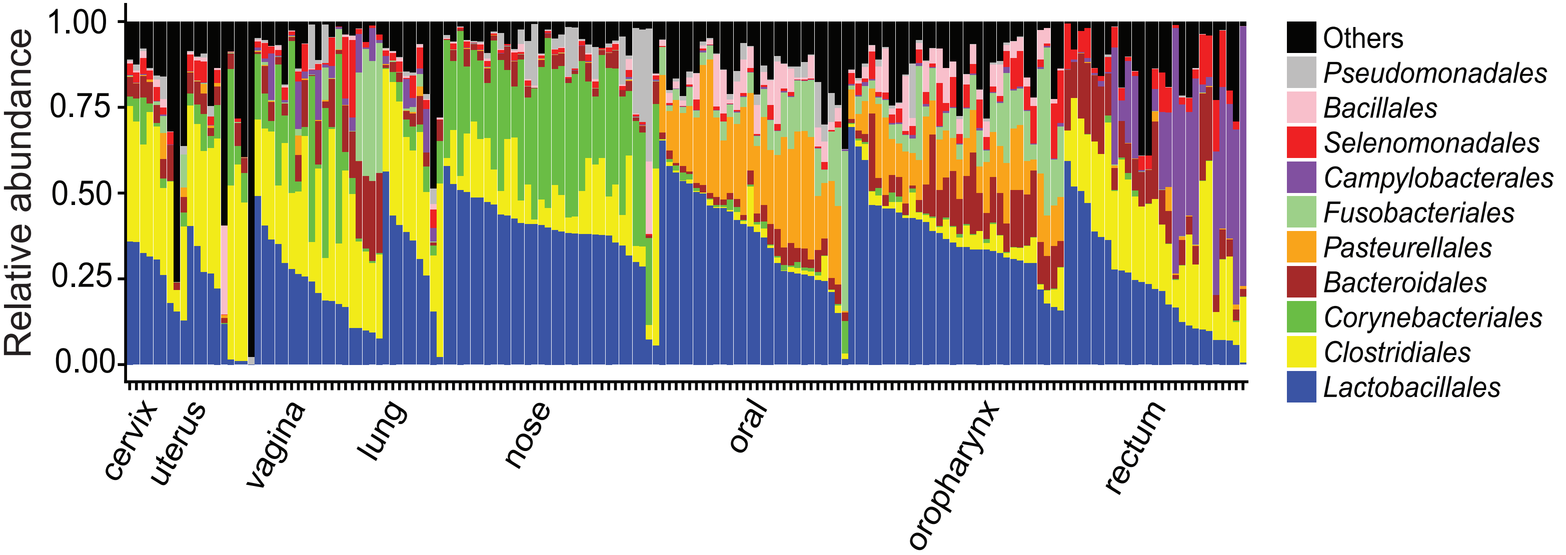
