## Supplementary material for "The microbiome of captive hamadryas baboon"

### Controls manage risk of contamination during wet-lab processing:

To avoid contamination risks, we strictly control the sampling process, DNA extraction, PCR and sequencing. All DNA extractions strictly followed the aseptic operation process under a clean bench. We also have DNA extraction negative control (from the DNA extraction to the sequencing process), sequencing blank control (clean water for sequencing), and sequencing positive control (mock community, *E.coli*).

As a result,

**1.** The total reads for sequencing blank control are only 61 and was then filtered.

**2.** The sequencing positive controls have highly similar microbial profiles with theoretical composition, as shown in the following figure. 99.95% of sequenced reads in the *E.coli* sample were assigned into *E.coli*. The bacterial profiles of mock are very similar with the expected composition.

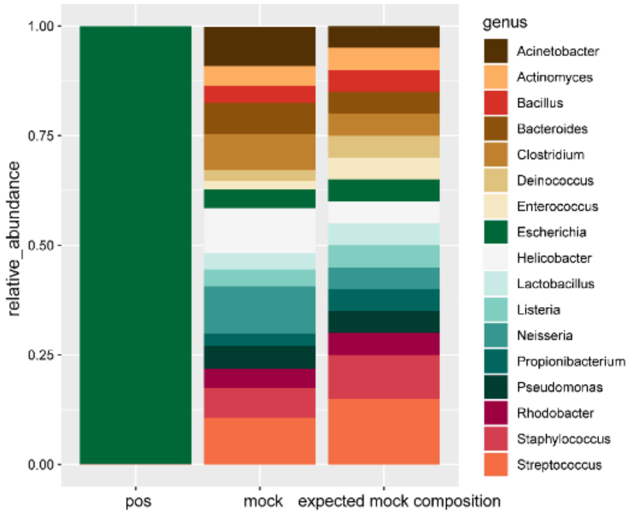


3. As Figure S4 of the manuscript showed, real samples are also different from DNA extraction negative control. The average unweighted unifrac distances between samples are shown as follows: the columns represent all the samples, yellow line represents mock control, red line represents DNA-extraction negative control, purple line represents *E.coli* positive control. The mock control, DNA extraction negative control and *E.coli* positive control have higher average UniFrac distances to other samples than most of the samples.

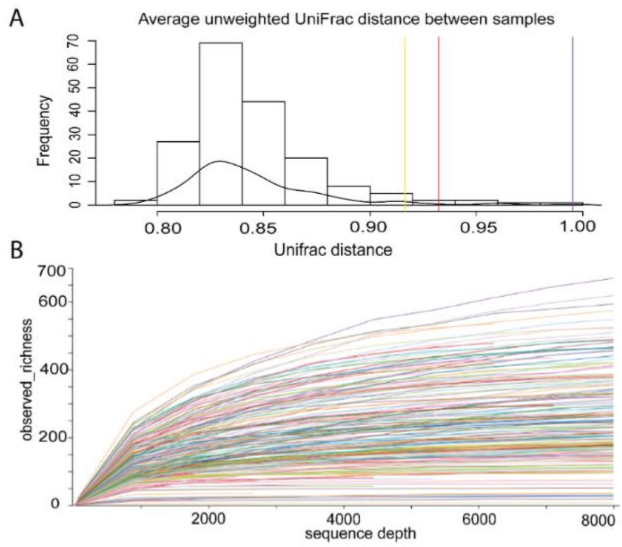


**<u>Sterile surgical procedure manage risk of contamination at sampling</u>**:

"*Animals were anesthetized for a full medical evaluation and physical examination.*" Importance: the animals being anesthetized allowed us to sample each site with extreme precision, as animals were not alert and did not move/shift during the sampling process.

"*Non-invasive samples (vagina, nose, oral, oropharynx, and rectum) were collected in a sheltered housing facility using sterile polyester swabs* (300263DNA, *Deltalab, Spain*)." Importance: conducting the sampling procedure in a sheltered facility and using new sterile swabs for each site further minimized possible contamination risks, eliminated the risk of environmental debris, and further added to the confidence of the swabs obtained. The swabs are DNAse and RNAse free, sterilized by ethylene oxide, and supplied in a polypropylene tube, which protects the sample prior to its analysis.

"*Following thorough medical evaluations, 8 of the animals were euthanized by a licensed veterinarian and a thorough postmortem examination was conducted in a separate necropsy room. Carcasses were opened ventrally to expose the organs, and all invasive sampling was performed sequentially from cranial to caudal.*" Importance: the autopsy and invasive sampling was conducted in a separate necropsy room. In addition to which, the organs were sampled sequentially from head to tail – samples from the upper organs were obtained before the lower organs. This anatomical direction of sampling eliminates the possibility of cross contamination where bacteria from the lower organs may be introduced into the upper organs, such as the possibility of transferring bacteria from the vagina/cervix/uterus to the lungs.

"*Lungs were excised, and both the left and right main bronchi were swabbed, whereas for all other invasive samples, a small surgical incision was made to obtain the swabs. A new sterile scalpel was used for each organ; new gloves were donned and new surgical utensils were used for each of the carcasses.*" Importance: these precautions further ensured that neither the surgical instruments nor the gloves would introduce any microbes into the organs/carcasses sampled sequentially.

As for the personnel that carried out the sampling procedures: Mads Frost Bertelsen is a licensed veterinarian at the Copenhagen Zoo and adjunct professor in veterinary clinical microbiology; he has performed sampling procedures such as these for a number of other peer-reviewed studies. Whereas Urvish Trivedi stems from a medical research background from the Dept. of Surgery, having experience in working with both animal models and patients that were part of peer-reviewed clinical studies.
